# Humans optimally anticipate and compensate for an uneven step during walking

**DOI:** 10.1101/2020.12.01.407023

**Authors:** Osman Darici, Arthur D. Kuo

## Abstract

The simple task of walking up a sidewalk curb is actually a dynamic prediction task. The curb is a disturbance that causes a loss of momentum, to be anticipated and compensated for. For example, the compensation might regain momentum and ensure undisturbed time of arrival. But without a selection criterion, there are infinite possible strategies. Here we show that humans compensate with an anticipatory pattern of forward speed adjustments, with a criterion of minimizing mechanical energy input. This is predicted by optimal control for a simple model of walking dynamics, with each leg’s push-off work as input. Optimization predicts a tri-phasic trajectory of speed (and thus momentum) adjustments, including an anticipatory, feedforward phase. In experiment, human subjects successfully regain time relative to undisturbed walking, with the predicted tri-phasic trajectory. They also scale the pattern with up- or down-steps, and inversely with average speed, as also predicted by model. Humans can reason about the dynamics of walking to plan anticipatory and economical control, even with a sidewalk curb in the way.

## INTRODUCTION

There are indeterminate control choices to be made during walking, not least when steady gait is interrupted by a surface perturbation such as a sidewalk curb. One solution is to do nothing until gait has actually been disrupted, and to rely on feedback control to restore stasis, similar to stabilizing standing balance after a perturbation. But an alternative possibility is to plan and act ahead with anticipatory, feedforward adjustments. For example, a ball, once sighted, may be caught by predicting its dynamics and planning and executing an interception course. A sidewalk curb may similarly be intercepted, albeit with most of the dynamics within the person rather than the object, and with a less clearly defined objective. This raises the question of what criteria govern the interception, whether by feedback or feedforward. If an objective advantage could be identified, it might also be sufficient to predict a single, optimal response. Thus, the seemingly simple task of dealing with an uneven step may yield insight on whether humans perform predictive, dynamical planning while they walk.

Both feedback and feedforward control could contribute to walking. Feedback control is important for balance during walking, for example to adjust foot placement in response to perturbations (Bauby and Kuo, 2000; O’Connor and Kuo, 2009; Wang and Srinivasan, 2014). In both standing and walking, feedback could be regarded as a means to control the body’s dynamical *state* (Kuo, 1995; Park et al., 2004). In contrast, feedforward is clearly used to plan the body’s location, for example to negotiate around obstacles or through doorways (Arechavaleta et al.; Brown et al., 2020). Motion must also be planned, with the help of vision, to step over upcoming obstacles (Patla, 1998), and to adjust foot placement (Matthis and Fajen, 2013). Thus, feedforward could be considered as planning of the body’s path, but not necessarily its dynamical state. But there may also be advantages to planning state, particularly speed or momentum. After all, it may help to speed up before jumping over a puddle. Indeed, runners do load the leg differently just before a drop (Müller et al., 2012), perhaps as a way to regulate momentum. The negotiation of uneven terrain might therefore benefit from feedforward planning of state. However, there currently lacks a mechanistic explanation or prediction for such planning.

Any systematic control strategy, regardless if feedforward or feedback, should also be driven by objective criteria to select among infinite options. An example is metabolic energy economy, which determines the preferred step length and step width of steady walking, as governed by the pendulum-like dynamics of the legs. Perhaps similar dynamics and similar economy apply to walking over uneven terrain as well. We previously explored this question with a simple model of human walking (Kuo, 2002), for the task of negotiating a single uneven step (termed Up-step or Down-step) during otherwise steady walking (Darici et al., 2020). An uncompensated Up-step would normally cause a loss of momentum, and thus a loss of time compared to walking the same distance uninterrupted. We used optimal control to determine the most economical strategy to regain lost time and momentum. The objective was to minimize a crude indicator of energy expenditure, the mechanical work performed in the step-to-step transition from one pendulum-like stance leg to the next (Donelan et al., 2002; Kuo et al., 2005). This yielded a strategy for negotiating an Up-step by modulating forward momentum over multiple steps, starting with a speed-up before the perturbation, and continuing the modulation for several steps after the perturbation. The model predicted a substantial economic advantage to the anticipatory speed-up, compared to post-hoc compensation alone. The optimal control was highly systematic, with an almost opposite strategy for negotiating a Down-step. This suggests that humans might also gain advantage from a feedforward, anticipatory strategy, which could allow a perturbation to be negotiated with no loss of time and good economy.

The criteria by which humans compensate for ground perturbations is unknown, as is whether they use feedforward or feedback. And even if energy expenditure were a concern, it might have minor import relative to other conceivable criteria. It is also quite possible that humans do not care about minor perturbations, and simply lose momentum to an Up-step. And even if momentum were regained through feedback control, they might still lose considerable time to the perturbation. We therefore sought to determine whether humans regulate their momentum, whether they can also regain lost time, and whether they use feedforward control in their compensatory strategy. The present study was therefore to experimentally test whether humans optimally compensate for a perturbation to step height and determine how that compensation is performed.

## METHODS

### Model of walking

We summarize predictions from an optimal control model of walking (Fig. 1; Darici et al., 2020), with details in Appendix. The task is to walk down a walkway interrupted by a single Up- or Down-step (numbered step *i* = 0; Fig. 1a), with adjustments to the forward speed *v*_*i*_ of each step. The model has rigid, pendulum-like legs supporting a point-mass pelvis of mass *M* (Fig. 1b; Kuo, 2002). The dynamics of the single stance phase are those of a simple inverted pendulum, which conserves mechanical energy and therefore loses speed stepping on an Up-step. As a simplification, the previous model’s (Kuo, 2002) swing phase dynamics are ignored, and the legs are constrained to fixed step lengths, similar to the “rimless wheel” model (McGeer, 1990). A level, nominal step (Fig. 1c) is punctuated by a step-to-step transition, where the trailing leg pushes off (PO) impulsively just before the leading leg’s dissipative collision (CO) impulse. This redirects the center-of-mass (COM) velocity to a new pendular arc described by the leading leg. The push-off and collision impulses are performed along the axis of the corresponding legs, with push-off as the only powered actuation, and (perfectly inelastic) collision the only dissipation. Experiments show that it explains how mechanical work and human metabolic energy expenditure increase as a function of step length (Donelan et al., 2002) or step width (in 3D model; Donelan et al., 2001) on level ground. Here we modeled uneven terrain as a small, vertical height discrepancy *b* in step height, where additional push-off can help compensate for momentum lost to an Up-step (Fig. 1d), and collision for momentum gained to a Down-step (Fig. 1e).

**Figure 1.**
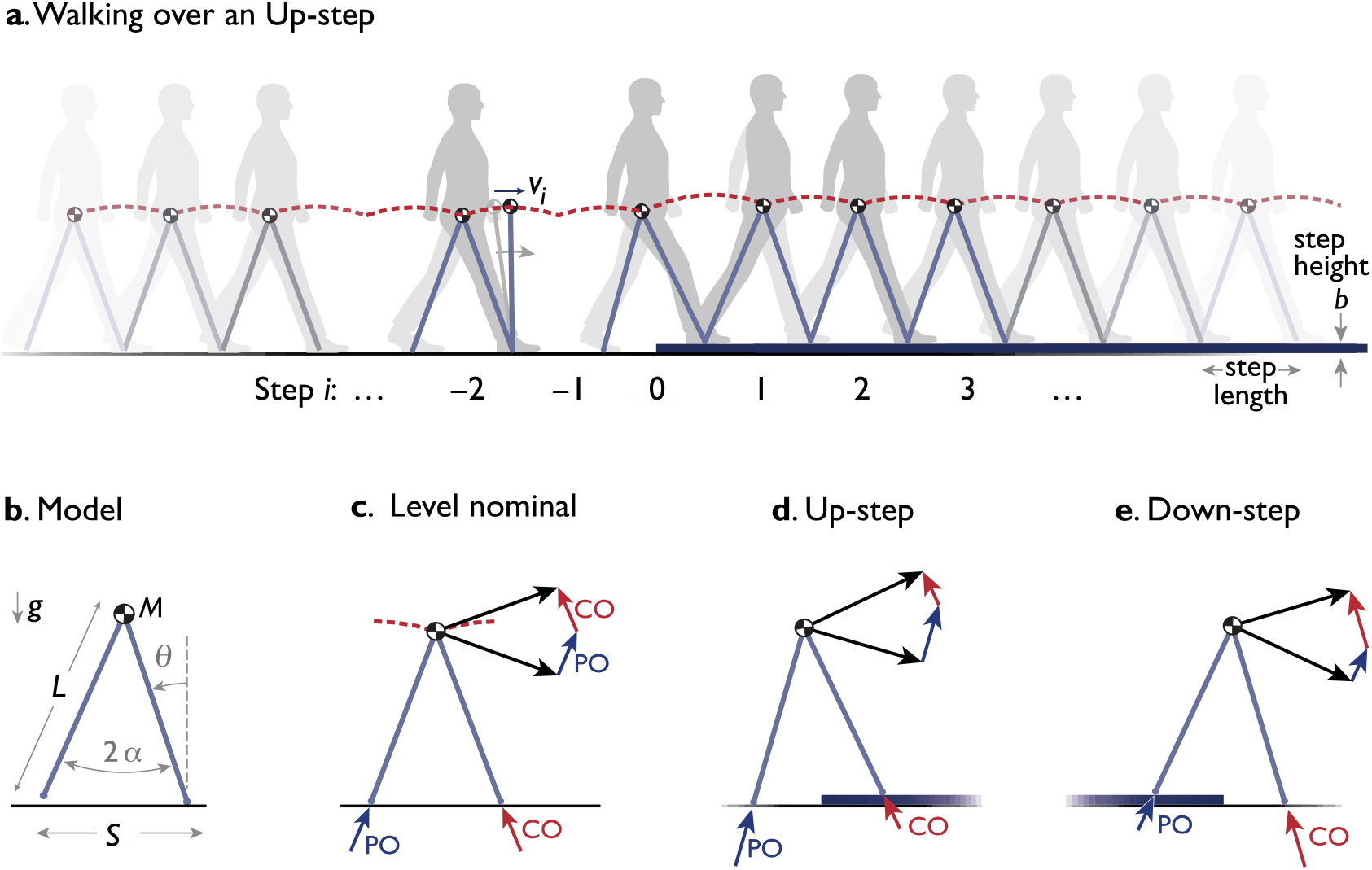
Dynamics of walking over a single Up- (or Down-) step. (**a**) Humans walk with the body center of mass (COM) moving up and down atop the stance leg behaving like an inverted pendulum. Momentum fluctuates in each step (numbered *i*), and is particularly disrupted by an uneven step (at *i* = 0). In experiment, human subjects walked in a walkway (30 m long) with level ground or a single Up- or Down-step (height *b* = 7.5 cm) at mid-point. Subjects were asked to walk this distance in roughly similar time, regardless of the perturbation, and without receiving feedback about time. Outcomes were quantified by the trajectory of speed fluctuations *v*_*i*_ at the discrete mid-stance instance (for 15 steps). (**b**) Dynamic walking model has a point mass *M* at pelvis, supported by an inverted pendulum stance leg (massless, length *L*, gravitational acceleration *g*, fixed step length *S*, fixed inter-leg angle 2*α*). (**c**) Level nominal walking has step-to-step transition where COM velocity (dark arrow) is redirected from forward-and-downward to forward-and-upward by an active, impulsive trailing leg push-off (PO), immediately before an inelastic, impulsive, leading leg collision (CO). Both PO and CO are directed along the corresponding leg. (**d, e**) The model walks Up or Down a step by modulating the sequence of push-offs surrounding and including the uneven step.

We formulated an optimal control problem for compensating for a single step height change (Darici et al., 2020). The objective was to minimize total push off work for multiple (*N* compensatory) steps while compensating for the terrain unevenness. The model was governed by the walking dynamics and constrained to start and end its compensation at steady, nominal speed, with the *N* steps distributed equally before and after the Up-step. It was also constrained to match the total time for nominal level, steady walking, thus making up for time lost to the Up-step. The decision variables were the push-off work *u*_*i*_ for each step (where *i* = 0 for the push-off onto the Up-step), causing changes in the forward speed *v*_*i*_ of each step, discretely sampled at the mid-stance instance when the stance leg passed through vertical, prior to the step-to-step transition. The control policy refers to the push-off sequence *u*_*i*_ (including *i* over a range of steps), or equivalently the sequence of speeds *v*_*i*_.

### Model Predictions

Our model predicted that there is considerable advantage of anticipation (Fig. 2). Considerable momentum and time are lost to an uncompensated Up-step (Fig. 2a). In contrast, the optimal policy is to speed up in advance of the Upstep (thereby reducing the loss of momentum and time atop it), and then regain momentum afterwards (Fig. 2b; Darici et al., 2020). For stepping down Fig. 2c), the optimal policy is almost exactly the opposite: slow down in advance, gain speed and time atop the Down-step, and then slow down again toward nominal speed. These strategies are executed through modulation in push-of work (Fig. 2d), with a peak in work when stepping onto the Up-step (and a minimum for Down-step). As a result, time (Fig. 2e) is first gained prior to the Up-step (and lost prior to Down-step), such that the cumulative time gain eventually reaches zero.

**Figure 2.**
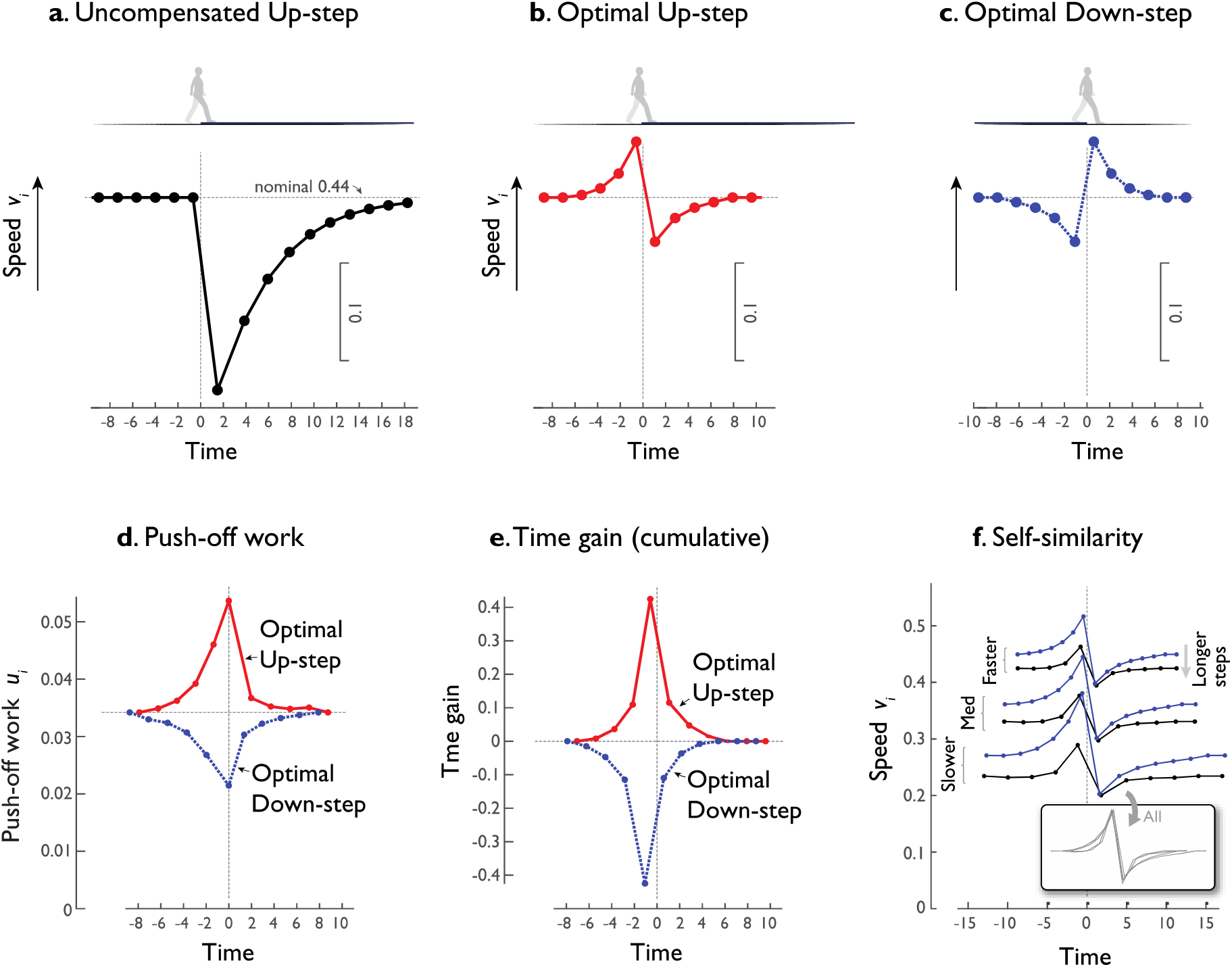
Model predictions for walking over an Up-step or Down-step. (**a**) Walking speed fluctuations vs. time, for level walking interrupted by a single Up-step (at time 0 and step *i* = 0) with no compensation (constant push-off), resulting in loss of momentum and time. Speeds *v*_*i*_ are sampled at mid-stance of each step (prior to step-to-step transition), and denoted by filled symbols. **(b)** Speed fluctuations for optimal Up-step compensation that minimizes push-off work. Model anticipates the perturbation with feedforward adjustment to speed up ahead of time, then loses momentum atop the perturbation, and then regains speed thereafter. **(c)** Speed fluctuations for optimal Down-step compensation (blue symbols) is nearly opposite in sign to the Up-step compensation (red): Slow down in advance, gain momentum, then slow down again. (**d**) Optimal control inputs are sequence of push-off work, shown for Up- and Down-step. Up-step requires more work, and Down-step less work, to walk same distance in same time. (**e**) Cumulative time gained for Up- and Down-step compensations. (f.) Self-similarity of Up-step compensations for three different nominal speeds and two different step lengths. All trajectories (see inset) are also scaled and super-imposed to illustrate self-similarity. For model predictions, conditions are similar to a human walking at 1.5 m/s with a 7.5 cm Up-step (nominal mid-stance velocity *V* = 0.44*g*^−0.5^*L*^0.5^, *S* = 0.79 *L*, *b* = 0.075*L*); described in detail by Darici et al. (2020).

An interesting feature of the optimal policy is self-similarity with respect to overall walking speed and step length (Fig. 2e). The pattern remains almost the same, only scaling in amplitude and time for different overall walking speeds or step lengths. The amplitude of speed fluctuations also scales inversely with speed, meaning slightly smaller fluctuations for faster speeds. This is because a step of fixed height (and thus gravitational potential energy) has a relatively smaller effect on the greater momentum (and thus kinetic energy) of faster walking. In addition, the timing scales such that the optimal strategy would be elongated in time with shorter step lengths, but in about the same number of steps. Thus, we expect similar fluctuation patterns regardless of an individual’s self-selected speed and step length, and slightly smaller fluctuations for faster overall speeds.

### Experiment

We measured speed fluctuations as humans walked down a level walkway (about 30 m) with a single, raised step onto a second level of 7.5 cm (Fig. 1A). We tested healthy adult subjects (*N* = 12; 7 male, 5 female, all under 30 yrs age), whose steps and walking speed were measured with inertial measurement units (IMUs) on both feet. There were three conditions: Up-step, Down-step, and Control on level ground. Both Up- and Down-step used the same walkway except in opposite directions, and Control took place on level floor directly alongside the walkway. The raised section, commencing about halfway down, was assembled from fairly rigid, polystyrene insulation foam. In all conditions, subjects walked at comfortable speed from a start line through and past a finish line. Trials took place in alternating direction, with a brief delay for the subject to turn around and stand briefly before starting the next trial. There were at least five (and up to eight) trials of each condition, usually with Up- and Down-step alternating with each other, except with occasional Control conditions inserted at random and interrupting that pattern. Before data collection, subjects were given opportunity to try the conditions and gain familiarity with the walkway and the location of the Up-step. For brevity, all mentions of the Up-step apply equally to the Down-step, unless explicitly stated. We tested one sample group, with sample size (*N* = 12) selected for a statistical power of at least 0.8, based on previous data for anticipated means and standard deviations (Darici et al., 2015). The experiment was performed once and each test subject was tested once with each condition as a repeated measure.

The experiment was minimally governed, aside from instructing subjects which conditions to perform. To establish a subjective “normal” walking speed, and walking time, subjects first performed two to four Control trials at the beginning of the experiment. They were then encouraged to walk in roughly similar time throughout the experiment. But they were never received feedback of their timing, even though such data were recorded by stopwatch. This was in part to mimic the unconstrained nature of daily living, and because the model’s predictions do not depend on a particular speed. We thus expected a range of speeds across trials and across subjects. To help subjects to step onto the Up-step without stutter steps, there was a visual cue (a paper sticker) on the floor, approximately 5 m from the Up-step. Subjects were informed that they could use this cue to line up their steps for the Up-step, although they were not required to use it, and no trials were excluded even if there was a stutter step. Anecdotally, most subjects appeared to pay little attention to the sticker, especially after the first few trials.

We measured walking speed trajectories with Inertial Measurement Units (IMUs) on each foot (Rebula et al., 2013). Each foot’s trajectory was determined by integrating inertial data, subject to an assumption that each foot comes briefly to rest at each footfall, approximately in the middle of stance. We estimated stride length and time from the forward distance and time between an IMU’s footfalls, respectively. Each foot’s speed was sampled discretely at each footfall instant as stride length divided by stride time. Individual distances traveled by the two feet were corrected for integration drift so that they both agreed on overall distance, using linear de-trending. Walking speed was estimated from the interlaced data from the two feet, in a discrete sequence termed IMU speed, with each sample assigned to the preceding mid-stance instance. To focus on speed fluctuations, data were analyzed for a central, 8.5 m segment of the walkway, or about 15 steps surrounding the Up-step. To compare between trials, the time *t* = 0 was defined as the instant of the footfall onto the Up-step (or Down-step, or step next to it for Control). This yielded a trajectory of walking speed for each trial. Each subject’s trials within a condition were averaged at discrete step numbers, as were the times for those steps, to yield an individual’s average trajectory per condition. As a measure of self-similarity (across different speeds and different subjects), an individual’s trajectories were compared against the average trajectory for all individuals with Pearson correlation coefficient *ρ*, with significant correlation determined by one-sample t-test. As a test of the model, individual trajectories were also correlated against the model’s predicted trajectories with a Pearson correlation coefficient *ρ*, again with one-sample t-test for significant correlation. Finally, a linear regression was performed to test for fluctuation amplitudes scaling negatively with walking speed, as predicted by the model.

## Results

We first examine overall walking speeds, as a basis for comparing speed fluctuations. For the central segment of the walkway, the overall average self-selected speed was 1.38 m/s on level ground (±0.10 m/s s.d. across subjects). Each individual typically had a small amount of variation in self-selected speed between trials, with about 5% c.v., coefficient of variation across control trials. Subjects were thus fairly consistent in their own walking speed, despite receiving no feedback regarding walking durations or speeds.

We next examine fluctuations in speed within each trial of level Control walking (Fig. 3, top row). The speed fluctuations were small in magnitude and noise-like, with variability 0.031 ± 0.007 m/s (root-mean-square variation within trial, reported as mean ± s.d. across subjects), or about 2.2% c.v. These fluctuations exhibited a small amount of systematic behavior, as demonstrated by correlating each individual’s average Control trial against the average Control across all subjects. The correlation *ρ* was 0.47 ± 0.31 (*P* = 2.5e-04), suggesting a degree of non-random systematicity between subjects, albeit of small amplitude within the 2.2% fluctuations.

**Figure 3.**
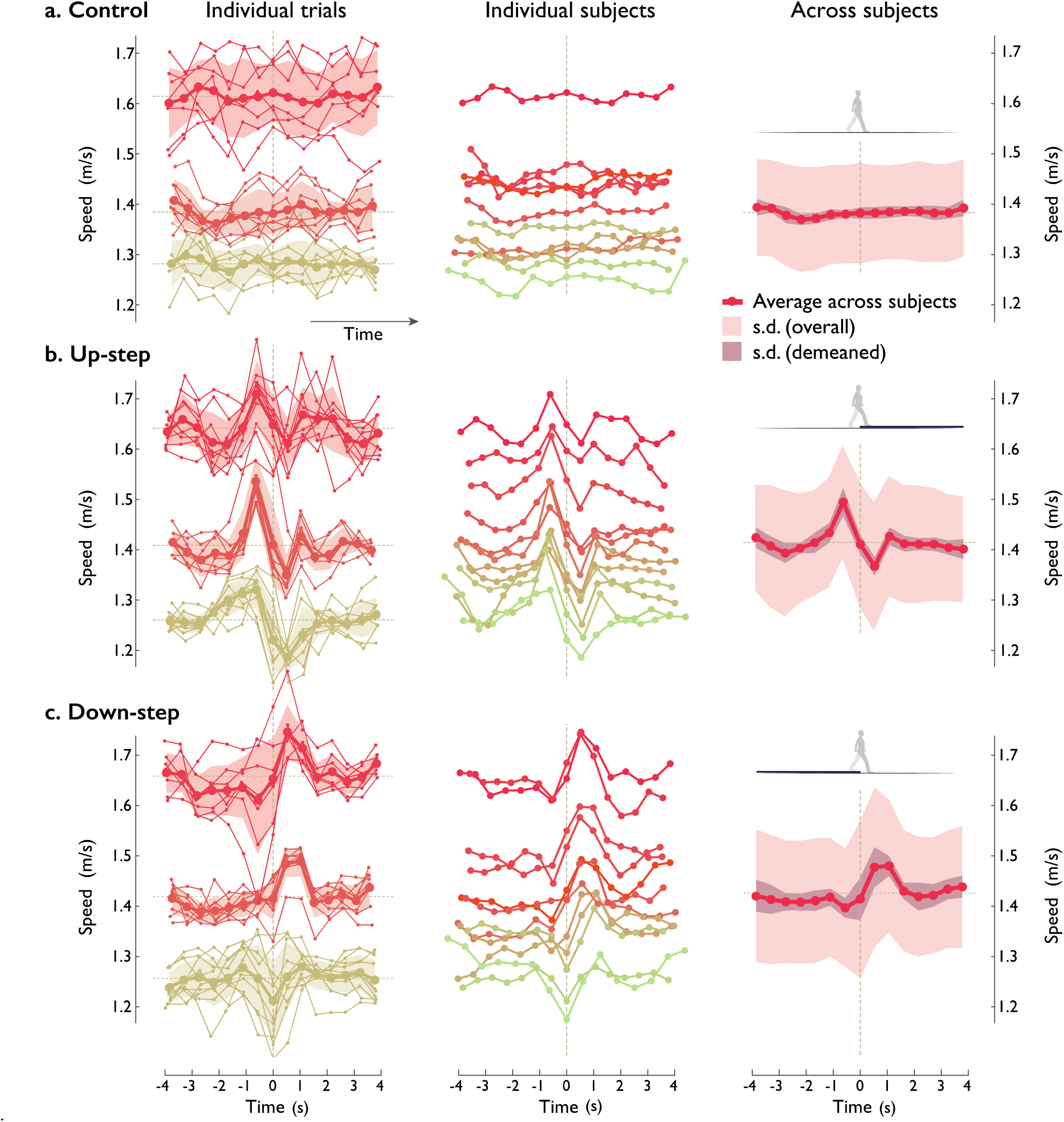
Human walking speed trajectories vs. time, for (A.) Control and (B.) Up- and (C.) Down-step conditions. Plots are arranged in columns: (left:) All individual trials of three representative test subjects (thin lines connecting small dots), along with per-subject average trajectories (across trials, thick lines) and standard deviations (shaded regions ±1 s.d.; dashed line indicates average speed). (Middle:) Speed fluctuations of each test subject (N = 12, individual colors) walking over the step, averaging all trials within each subject. (Right:) Average speed trajectories across all test subjects (solid line), with standard deviation across all subjects (light shaded region), and standard deviation ignoring subject-dependent speed (darker shaded region). Speed is defined as step length divided by step time, assigned to the middle-stance instant of each step (indicated by dot symbols). All trials are aligned to zero time, defined as middle-stance instant after landing on the Up- or Down-step (both 7.5 cm high).

### 1. Humans compensated for Up- and Down-steps to conserve walking duration

Subjects walked at similar overall speeds whether or not there was an uneven step. There were no significant differences in overall speed, step length, or segment duration due to condition (*P* = 0.65, *P* = 0.78, *P* = 0.96 respectively, repeated measures ANOVA). Speeds were also fairly consistent across trials within Up- or Down-step conditions (2 – 3% c.v.). This compensation contrasts with what would be expected for a no-compensation strategy. The model, if performing constant push-offs instead of compensating, would lose about 1 s on the Up-step compared to the Down-step (Fig. 2a), compared to level walking. Alternatively, a particle sliding on frictionless ground at human-like speed, would be expected to lose about 0.7 m/s and 8 s to an upward ramp of equivalent height. Thus, both a walking model and a sliding particle lose substantial speed and time due to a change in height, if not for some form of active compensation. In contrast, human subjects maintained almost the same walking speed and duration, as expected of successful compensation for an Up- or Down-step.

### 2. Up- and Down-step compensatory speed fluctuations were systematic and self-similar

There was also a clear pattern in compensations for an uneven step, with consistent fluctuations in walking speed (Fig. 3). The fluctuations within these trials were greater than those of Control, about 3.0% and 3.4% c.v. for Up- and Down-steps (Fig. 3 middle and bottom), respectively. The compensation strategies, in terms of walking speed trajectory over time, appeared qualitatively similar between multiple trials for an individual (Fig. 3 left column), and between different individuals (Fig. 3 middle column), to yield a single representative trajectory for all Up-step compensations (Fig. 3 right column).

The basic response could be summarized in terms of a triphasic pattern centered about the Up-step: (1) Speed up in the two steps prior, (2) then lose speed during the two steps onto the Up-step and immediately thereafter, and (3) then regain speed over the following one or two steps. The peak speed just prior to the Up-step (*i* = −1) was about 5.7% greater than average speed, and the minimum after the Up-step (*i* = 1) was about 3.4% slower. Similar observations were the case for Down-step compensations (Fig. 3 bottom row), except that fluctuations were in nearly the opposite direction, with a basic pattern of slow down, speed up, slow down. The timing was slightly different, with the initial slow-down being clearest for only one step immediately before the Down-step (*i* = −1), then speed-up occurring for about three steps, and the return to normal walking in only about one step.

The systematic behavior was quantified as follows. The Up-step and Down-step conditions were either not correlated or very weakly correlated with Control (*ρ* = −0.016 ± 0.21 and 0.184 ± 0.193, respectively, correlating each subject against Control average across subjects, *P* = 0.78, *P* = 0.007). But the patterns were similar between each individual’s Up-steps, with a positive correlation coefficient between Up-step trials (*ρ* = 0.82 ± 0.1252; correlating each subject against Up-step average across subjects, *P* = 1.26e-10 paired t-test of correlations). The same was true for Down-step patterns, with positive correlation (*ρ* = 0.68± 0.27, *P* = 3.0e-06). Moreover, the two fluctuations for the two conditions were somewhat opposite to each other, with a negative correlation between individual Up-steps and average Down-step pattern and vice versa (respectively *ρ* =−0.34 ± 0.16, *P* = 1.6e-05; *ρ* =−0.27 ± 0.22, *P* = 0.0016).

### 3. Step lengths and times also fluctuated

Step lengths and times also appeared to have fluctuation patterns (Fig. 4), reported descriptively here. For Up-steps (Fig. 4 left/top row), step lengths were about +3.1%, +18.5%, and −7% for the three steps surrounding the perturbation (*i* = −1, 0, +1), respectively, compared to overall walking speed. For Down-steps, the corresponding step length differences were .24% 2.29% −2.58% (Fig. 4 bottom row). As for step times (Fig. 4 bottom), the Up-step differenceswere-5.8%, +15%, −2.7%, and Down-steps differences were +2.5%, +2.5%, and −3.5%, compared to average step period. No statistical tests were performed, as the model made no specific predictions about these fluctuations.

**Figure 4.**
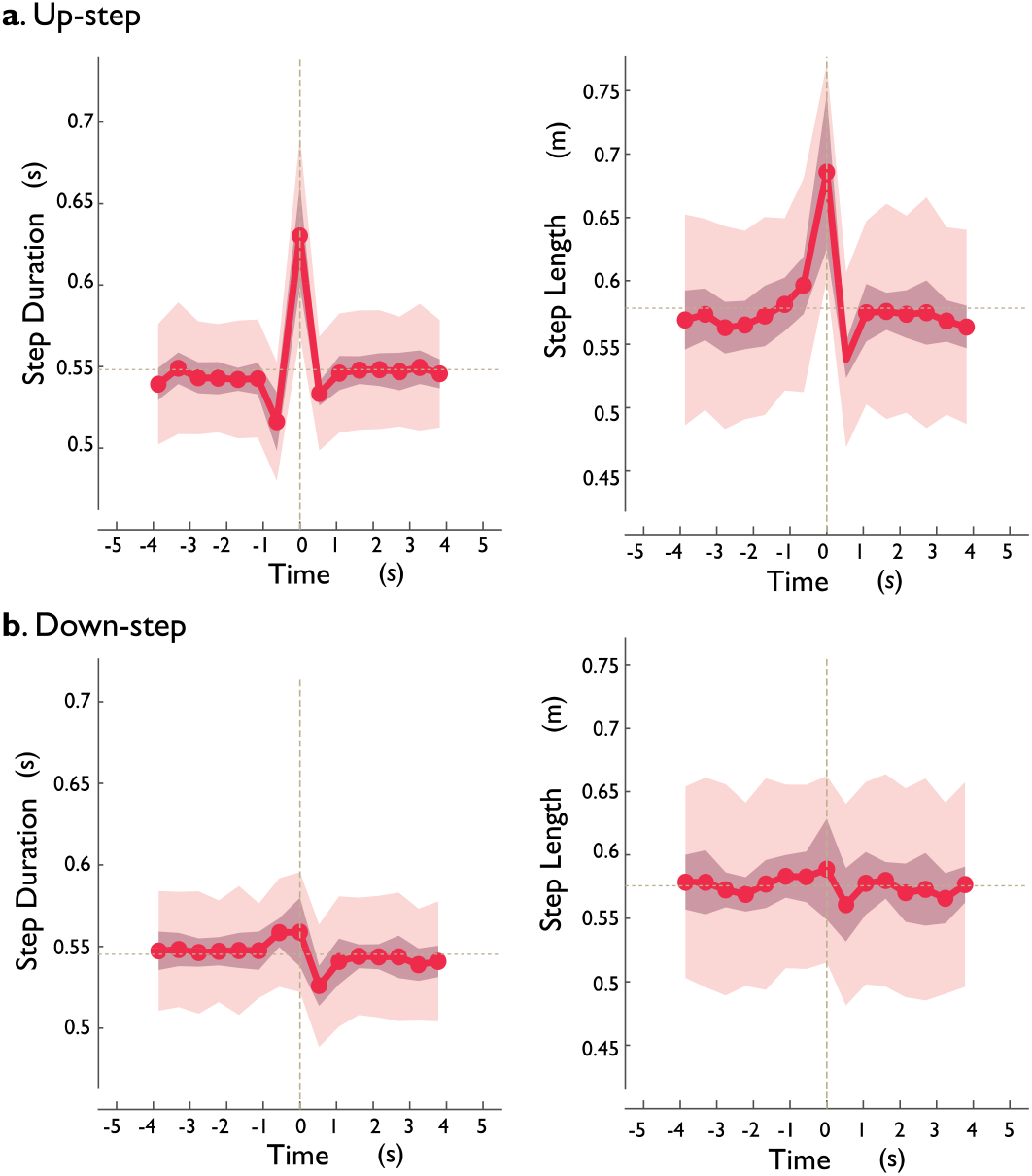
Human (left column:) step time and (right column:) step length fluctuations versus time, for (a.) Up-step and (b.) Down-step. Shown are step times and step lengths for each step (line denotes mean across subjects, shaded area denotes ±1 s.d.; *N* = 12) vs. time, with vertical line denoting the step onto the perturbation.

### 4. Optimization model predicts humans walking speed compensations

The human compensation strategies agreed reasonably well against model predictions (Fig. 5a, b for model; c, d for human). Both exhibited a general response of speeding up before the Up-step, then slowing down on and after, and finally regaining speed toward nominal speed. Down-step responses also agreed, with approximately opposite pattern to Up-step. This agreement was quantified by a positive correlation coefficient between human and model fluctuations for both Up-steps and Down-steps (*ρ* = 0.50 ± 0.21, *P* = 4.4e-6 and *ρ* =0.59 ± 0.17, *P* = 1.08e-7). And in keeping with the model’s prediction of opposing fluctuations for Up- vs. Down-steps, there was also a negative correlation between human Up-steps and model Down-steps, and vice-versa (*ρ* = −0.42 ± 0.21, *P* = 2.75e-5 and *ρ* =−0.54 ± 0.15, *P* = 8.05e-8). We also verified that human control responses were not correlated with model predictions for either Up- or Down-steps, with correlation coefficients not significantly different from zero (P = 0.32, P = 0.31).

**Figure 5.**
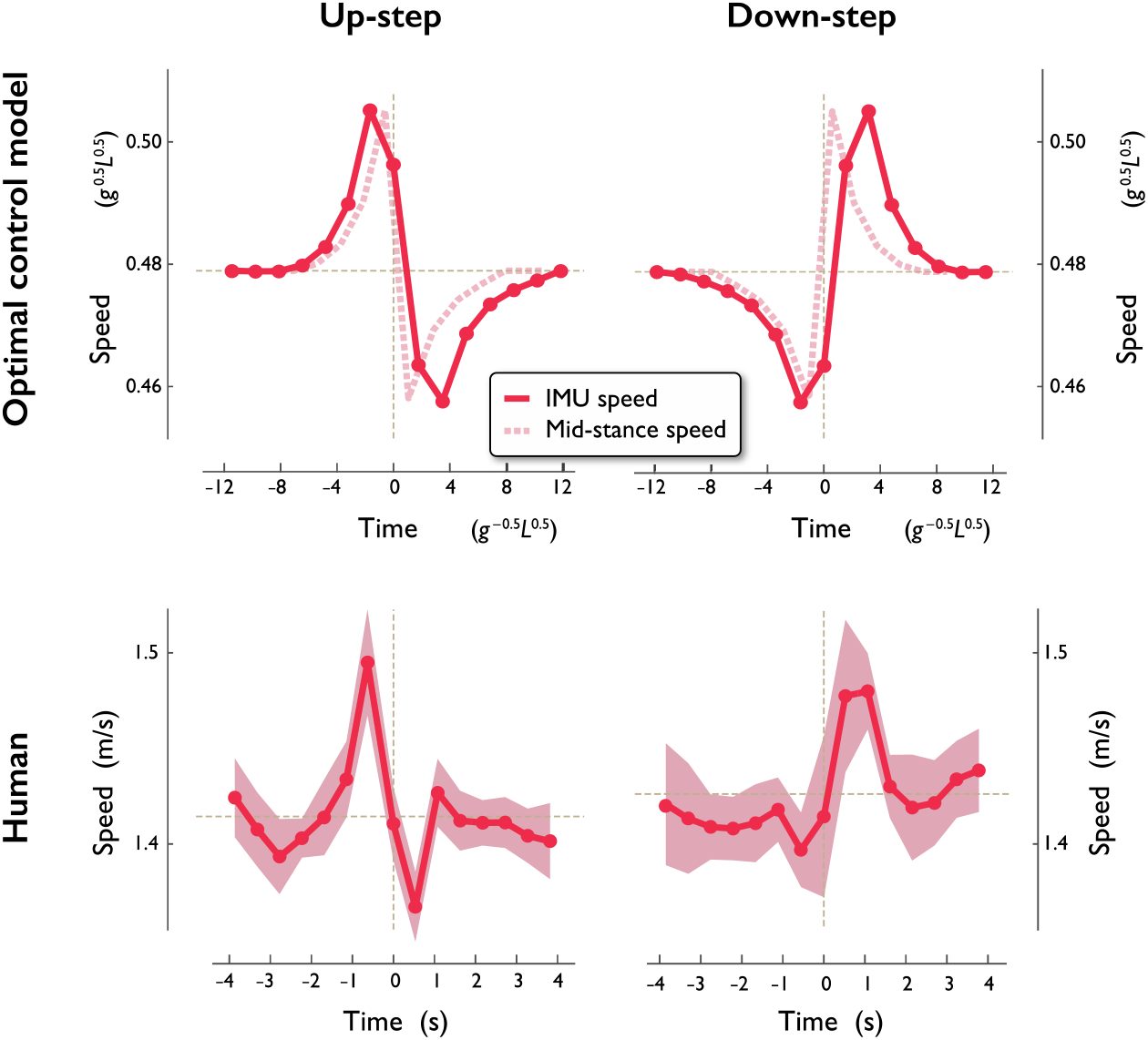
Comparison of model and human walking speed fluctuations vs. time, compensating for (left column:) Up- and (right column:) Down-steps. (Top row:) Model speed fluctuations predicted to minimize push-off mechanical work (Bottom row:) Experimentally measured compensation strategies for humans (N = 12), showing average speed pattern across subjects (shaded regions denote ±1 s.d. after eliminating variations in average speed). Each data point corresponds to speed measred by inertial measurement unit (IMU speed), defined as step distance divided by step time, and assigned to the instance when the stance leg is upright. The first step onto the Up- or Down-step is indicated by vertical dashed line, also at middle stance instant. The average walking speed is denoted by horizontal solid line. Model trajectories are converted from mid-stance speed for simulation into the equivalent experimental IMU speed, plotted in dimensionless speed and time, equivalent to units of 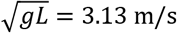 and 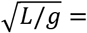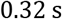, respectively (using gravitational acceleration *g* and human leg length *L* = 1 m).

Human responses also exhibited negative scaling with respect to walking speed, as predicted by model. A linear regression of normalized speed fluctuation amplitudes vs. overall speed revealed a negative coefficient (−1.64 ± 0.58 s/m^2^, mean ± c.i., *P* = 2.6e-12), meaning that a 1 m/s increment in overall speed was accompanied by an approximately 12.3% reduction in fluctuation magnitude for a 7.5 cm step.

## Discussion

We examined how humans anticipate and compensate for a step change in the height of an otherwise flat walking surface. The compensatory response was characterized by a systematic, tri-phasic pattern in walking speed fluctuations, from which we draw three notable observations. First, the response exhibited self-similarity, in that the same basic pattern could describe behavior at a variety of average walking speeds and step lengths. Second, the response also exhibited an anticipatory component, meaning that it partially occurred prior to physically encountering the step. Finally, the response was consistent with predictions from a simple walking model, optimizing for least mechanical work. We next discuss these findings with regard to implications for feedforward human control.

Human speed fluctuations were quite systematic. They exhibited a similar basic pattern across different trials of an individual, across different individuals, and at a variety of average walking speeds and step lengths (Figs. 3 & 4). Part of the systematicity could be attributed to dynamics, with a pendulum-like exchange of speed for height atop the Up-step (*i* = 0, Fig. 3), along with a loss of time (Fig. 4). But some of the systematicity is attributable to active control, because speed was then quickly regained toward nominal, considerably faster than expected if there were no active compensation (Fig. 2a; see also Darici et al., 2020). And more telling was the speed-up prior to the perturbation, which cannot be attributed to a feedback response to perturbation, but rather indicates intentional, anticipatory control. In addition, the speed fluctuations were lower in amplitude for higher speeds, as predicted for step-to-step transition dynamics. The compensatory strategies therefore reflect pendulum-like dynamics, and systematic, central nervous system control with feedback and feedforward (anticipatory) components.

The result of this active control was successful compensation for time lost to the perturbation. Overall walking duration was conserved across experimental conditions, demonstrating an ability to correct for uneven terrain. This cannot be explained by post hoc, feedback regulation of instantaneous speed, which would restore nominal speed but with a loss (gain) of time from the Up-step (Down-step). Nor can it be explained by learned adaptation during the experiment, because subjects never received feedback about their walking duration, and were only loosely advised to keep that time consistent. The control, particularly the anticipatory component, instead appears to be based on prior knowledge or experience. In daily living, humans regularly make decisions regarding walking route and speed, and seem able to estimate what path may take less time or effort. They may accumulate considerable experience, perhaps equivalent to an internal model of walking dynamics, sufficient to plan anticipatory compensations. Learning and anticipatory planning have mainly been addressed by the separate field of neuromotor control, which has theorized that upper extremity reaching movements are planned with CNS internal models of dynamics, and driven by an objective of movement accuracy (Franklin et al., 2008; Sharp et al., 2011). The present study borrows from that approach in its use of dynamical modeling and optimal control computations.

This compensatory strategy is consistent with a simple, optimal control model of walking. There are infinite ways to walk over an up-step perturbation without suffering a loss of time. But minimization of work for step-to-step transitions predicts the particular triphasic pattern observed here. A small (7.5 cm high) step might seem too trivial to compensate for, but our model suggests that considerable time could be lost (Darici et al., 2018), and substantial energy lost without anticipation (Darici et al., 2020). Humans seem well able to gauge a relatively small surface irregularity, plan a dynamical course of action, and then execute that plan for several steps before and after the perturbation. They appear capable of reasoning about the dynamics of walking.

This raises the question how the control is implemented by the central nervous system. The human’s ability to reason about surface perturbations could be regarded as equivalent to performing optimal control with an internal model of walking dynamics. But its neural representation need not resemble optimal control. For example, reinforcement learning suggests that an objective function such as ours could be optimized iteratively, and expressed as a function of body state and terrain, starting from a visual terrain image as input (Heess et al., 2017). The resulting control policy is a mapping from vision and state to action, which might be considered an inverse internal model of dynamics (Kawato, 1999). Our results raise the possibility that such a mapping could be simple and scalable. A single Up-step response could be learned, and then merely scaled in amplitude for other walking speeds or step heights, due to the systematic nature of the dynamics. Thus, the control policy might be stored in quite compact form, a possibility raised but yet to be tested. Also needed for learning is a means to evaluate the objective cost function. Our cost of mechanical work could be evaluated by body somatosensors, but information might also be gained from physiological sensors of metabolic cost. In fact, the work of step-to-step transitions exacts an approximately proportional metabolic cost in steady state walking (Adamczyk et al., 2006; Donelan et al., 2001; Donelan et al., 2002; Kuo et al., 2005). It remains to be tested whether the same holds true for the transient conditions examined here, but metabolic energy is compelling for its physiological relevance and importance for animal life (Alexander, 1996). Thus, the optimal compensation could be learned from feedback of physiologically relevant information.

This study highlights a less-appreciated aspect of vision-based path planning. It is clear that humans use vision to plan paths for the body (Arechavaleta et al.; Brown et al., 2020), including adjustment of COM height (Müller et al., 2012) and foot placement (Matthis and Fajen, 2013; Patla, 1998). But we found that humans plan not just positions, but also momentum. They make quick, dynamically sensible decisions to overcome quite minor obstacles, apparently for energetic benefit. Such planning might also explain the leg loading preceding a Down-step in human running (Müller et al., 2012). It is also consistent with how birds run over an obstacle, with an anticipatory vault in the step beforehand (*i* = −1), perhaps for economy (Birn-Jeffery et al., 2014). Path planning may therefore be for more than just body location, but also dynamical state, and for the purpose of energy economy.

The present model has a number of limitations. One is that Down-step responses were predicted less well than Up-steps (Fig. 5). We suspect that the inverted pendulum analogy is less predictive for stepping down, when humans may allow the trailing knee to flex, perhaps to reduce the rate of fall. Our model might be improved by inclusion of a knee (e.g., Dean and Kuo, 2009) and feet (Zelik et al., 2014), which would better reflect the fore-aft asymmetries of the human, active lifting of the foot when needed (Wu and Kuo, 2016), and perhaps predict the asymmetries observed in Up- vs. Down-steps. We also modeled only fixed step lengths, but inclusion of variable step lengths or foot placements (Bhounsule, 2014; Kuo, 2001; Ojeda et al., 2015) might help to predict the variations in step length observed experimentally (Fig. 4). It is also possible that humans couple their sagittal and frontal plane motions for a change in step height, which might be accommodated in a three-dimensional model (Kim and Collins, 2017; Kuo, 1999). Additional degrees of freedom might help to predict the multi-joint actions of humans, given hypotheses regarding the attendant costs. As a simplification, we also examined only single terrain disturbances. But we consider it relatively straightforward to predict and test compensation strategies for more complex terrain disturbances over multiple steps. Fortunately, the dynamical modeling approach is amenable to inclusion of more degrees of freedom, and to more rigorous experimental testing.

Another limitation was that the human speed fluctuations were simply noisy. There was significant variability between trials of a single individual and between different individuals. This was due in part to the relatively small step height perturbations, which resulted in relatively small speed fluctuations compared to the noisy intrinsic variability of humans. We intentionally selected small perturbations to remain within the realm of pendulum-like walking. We would expect relatively less noise with larger step height changes, which would likely necessitate more human-like features in the model, such as the knees mentioned above.

Despite these limitations, we showed here that humans perform anticipatory speed adjustments on uneven terrain. A simple model minimizing the mechanical work of step-to-step transitions can predict these adjustments. The adjustments start several steps before, extend after the perturbation, in a tri-phasic pattern of speed fluctuations. Such a pattern is consistent with metabolic energy expenditure as a criterion for optimal control and shows that humans perform feedforward control before a perturbation is directly encountered. The central nervous system appears to anticipate the effects of disturbances on the dynamics of the body and exploit these dynamics for active and economical control.

## Acknowledgements

This work is supported by NSF DGE 0718128, the ONR ETOWL program, NIH AG030815, the Dr. Benno Nigg Research Chair (University of Calgary), and NSERC (Natural Sciences and Engineering Research Council of Canada) Discovery program and Canada Research Chair (Tier 1) program.

## Appendix

## Dynamic walking model

The model dynamics are briefly summarized as follows (detailed previously by Darici et al., 2018). Each of *N* steps has index *i* with the Up- or Down-step disturbance located at *i* = 0 (Fig. 1c). Negative *i* therefore refer to the preparatory steps beforehand, and positive to recovery steps thereafter. Each step has a pendulum-like single stance phase with passive dynamics, and a costly step-to-step transition. Mechanical work is only performed during that transition, starting with COM velocity 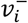 directed forward and downward at the end of each stance phase. For brevity, the equations presented here use dimensionless versions of quantities, with *M*, *g*, and *L* as base units. The step-to-step transition starts with pre-emptive push-off work *u*_*i*_ (in units of mass-normalized work) performed impulsively along the trailing leg to redirect the COM velocity. This is followed immediately by the heel-strike collision along the leading leg, to yield post-collision velocity 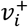. Again applying impulse-momentum (Kuo, 2002),

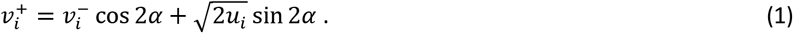

Another single stance phase follows the step-to-step transition, and is modeled as an underactuated, simple inverted pendulum. As a discrete indicator of overall forward momentum, we use the mid-stance velocity *v*_*i*_ (no superscript; see Fig. 1) following step-to-step transition 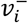, sampled when the leg is vertical and the COM velocity is purely forward.

We treat steady, level walking as the nominal condition (Fig. 1c). The nominal push-off work *u*_*i*_ offsets the collision work (Kuo, 2002), so that

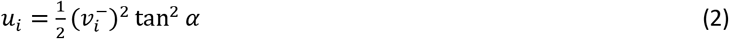

and 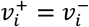. The uneven step disturbs steady walking (Fig. 1d). Its height *b* (positive for up-steps, negative for down-steps) causes the preceding stance phase to end with a different stance leg angle from nominal. For a given height *b* and step length *S*, we define the angular disturbance as *δ*_*i*_,

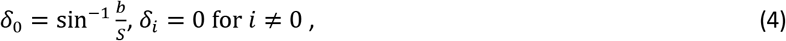

where the angle is zero for all non-disturbance steps.

An inverted pendulum stance phase follows each step-to-step transition. A step time *τ*_*i*_ defined as the time for the stance leg angle *θ* to move between successive step-to-step transitions, from 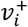 to 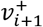 and passing through mid-stance speed *v*_*i*_.

Using the linearized dynamics, the dimensionless step time *τ*_*i*_ of step *i* is

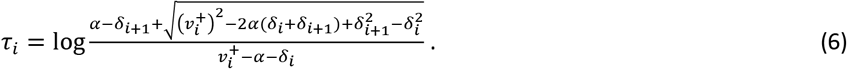

Solving the equation of motion with the step time, the velocity at end of stance 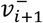, or equivalently the beginning of the next step-to-step transition can be found as:

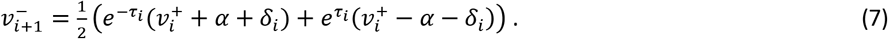

Mid-stance time 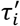 for step *i* can also be found using the linearized dynamics:

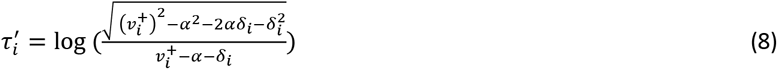

Solving for mid-stance speed *v*_*i*_,

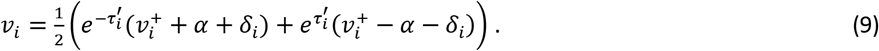

We chose nominal parameters to correspond to typical human walking. A person with leg length *L* of 1 m may typically walk at 1.5 m/s, with step length of 0.79 m and step time of 0.53 s (from anecdotal observations). Using dynamic similarity, parameters and results may be expressed in terms of body mass *M*, gravitational acceleration *g*, and *L* as base units. The corresponding model parameters treated as dimensional are angle *α* = 0.41, push-off *U* = 0.0342 *M*_*g*_*L*, step time *T* = 1.665 *g*^−0.5^*L*^0.5^, and pre-collision speed *V*^∗^ = 0.601 *g*^0.5^*L*^0.5^, where capital letters indicate nominal values for *u*_*i*_, *τ*_*i*_, and 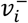, respectively. We also refer to a nominal speed *V* = 0.44 *g*^0.5^*L*^0.5^ for midstance speed *v*_*i*_. We considered a range of up-step heights, for example *b* = 0.075*L*, equivalent to about 7.5 cm for a human.

## Optimization problem

The optimization is formulated as follows, with policy *π* denoting the set of push-offs *u*_*i*_:

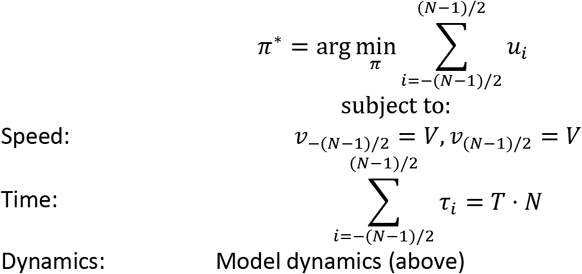

where *N* is the total (odd) number of steps and step *i* = 0 is the first step on the Up-/Down-step. Thus *N* adjusts how far in advance and or after the perturbation for which the model can modulate its momentum or speed. The speed constraints are such that the initial and final conditions are equal to the nominal, steady speed *V*. The time constraint makes up for lost time, so that the total time is equal to the nominal time to walk *N* steps on level ground. By the end of the control sequence, the model must walk at the same speed as nominal and must have caught up with the nominal model on level ground. We chose *N* large enough to cover the speed adjustments that humans made in the experiments. Note that, because human speeds are most conveniently measured from footfall to foot-fall, we converted the model speeds to a similar footfall definition (stride length divided by stride time, footfall to footfall) for purposes of comparison between model and human (Fig. 5).

## References

Adamczyk, P. G., Collins, S. H. and Kuo, A. D. (2006). The advantages of a rolling foot in human walking. Journal of Experimental Biology 209, 3953–3963.

Alexander, R. M. (1996). Optima for Animals. Princeton, NJ: Princeton University Press.

Arechavaleta, G., Laumond, J., Member, S., Hicheur, H. and Berthoz, A. An optimality principle governing human walking. Robotics, IEEE Transactions on 1–5.

Bauby, C. E. and Kuo, A. D. (2000). Active control of lateral balance in human walking. J Biomech 33, 1433–1440.

Bhounsule, P. (2014). Control of a compass gait walker based on energy regulation using ankle push-off and foot placement. Robotica 33, 1–11.

Birn-Jeffery, A. V., Hubicki, C. M, Blum, Y., Renjewski, D., Hurst, J. W. and Daley, M. A. (2014). Don’t break a leg: running birds from quail to ostrich prioritise leg safety and economy on uneven terrain. Journal of Experimental Biology 217, 3786–3796.

Brown, G. L., Seethapathi, N. and Srinivasan, M. (2020). Energy optimality predicts curvilinear locomotion.

Darici, O., Arthur D., K. and Hakan Temeltas (2015). Beware of the bump: Optimal strategy to traverse a step height perturbation. In Dynamic Walking, p. Ohio, USA.

Darici, O., Temeltas, H. and Kuo, A. D. (2018). Optimal regulation of bipedal walking speed despite an unexpected bump in the road. PLOS ONE 13, e0204205.

Darici, O., Temeltas, H. and Kuo, A. D. (2020). Anticipatory Control of Momentum for Bipedal Walking on Uneven Terrain. Scientific Reports 10, 540.

Dean, J. C. and Kuo, A. D. (2009). Elastic coupling of limb joints enables faster bipedal walking. J R Soc Interface 6, 561–573.

Donelan, J. M., Kram, R. and Kuo, A. D. (2001). Mechanical and metabolic determinants of the preferred step width in human walking. Proc. Biol. Sci 268, 1985–1992.

Donelan, J. M., Kram, R. and Kuo, A. D. (2002). Mechanical work for step-to-step transitions is a major determinant of the metabolic cost of human walking. Journal of Experimental Biology 205, 3717–27.

Franklin, D. W., Burdet, E., Peng Tee, K., Osu, R., Chew, C.-M., Milner, T. E. and Kawato, M. (2008). CNS Learns Stable, Accurate, and Efficient Movements Using a Simple Algorithm. J Neurosci 28, 11165–11173.

Heess, N., Tb, D., Sriram, S., Lemmon, J., Merel, J., Wayne, G., Tassa, Y., Erez, T., Wang, Z., Eslami, S. M A., et al. (2017). Emergence of Locomotion Behaviours in Rich Environments. arXiv:1707.02286v2.

Kawato, M. (1999). Internal models for motor control and trajectory planning. Current Opinion in Neurobiology 9, 718–727.

Kim, M. and Collins, S. H. (2017). Once-Per-Step Control of Ankle Push-Off Work Improves Balance in a Three-Dimensional Simulation of Bipedal Walking. IEEE Transactions on Robotics 33, 406–418.

Kuo, A. D. (1995). An optimal control model for analyzing human postural balance. IEEE Trans Biomed Eng 42, 87–101.

Kuo, A. D. (1999). Stabilization of lateral motion in passive dynamic walking. International Journal of Robotics Research 18, 917–930.

Kuo, A. D. (2001). A simple model of bipedal walking predicts the preferred speed-step length relationship. Journal of Biomechanical Engineering 123, 264–9.

Kuo, A. D. (2002). Energetics of actively powered locomotion using the simplest walking model. Journal of Biomechanical Engineering 124, 113–20.

Kuo, A. D, Donelan, J. M. and Ruina, A. (2005). Energetic consequences of walking like an inverted pendulum: step-to-step transitions. Exercise and sport sciences reviews 33, 88.

Matthis, J. S. and Fajen, B. R. (2013). Humans exploit the biomechanics of bipedal gait during visually guided walking over complex terrain. Proc. Biol. Sci. 280, 20130700.

McGeer, T. (1990). Passive dynamic walking. International Journal of Robotics Research 9, 62–82.

Müller, R., Ernst, M. and Blickhan, R. (2012). Leg adjustments during running across visible and camouflaged incidental changes in ground level. Journal of Experimental Biology 215, 3072–3079.

O’Connor, S. M. and Kuo, A. D. (2009). Direction-dependent control of balance during walking and standing. J. Neurophysiol 102, 1411–1419.

Ojeda, L. V., Rebula, J. R., Kuo, A. D. and Adamczyk, P. G. (2015). Influence of contextual task constraints on preferred stride parameters and their variabilities during human walking. Medical Engineering & Physics 37, 929–936.

Park, S., Horak, F. B. and Kuo, A. D. (2004). Postural feedback responses scale with biomechanical constraints in human standing. Exp Brain Res 154, 417–427.

Patla, A. E. (1998). How is human gait controlled by vision. Ecological Psychology 10, 287–302.

Rebula, J. R., Ojeda, L. V., Adamczyk, P. G. and Kuo, A. D. (2013). Measurement of foot placement and its variability with inertial sensors. Gait Posture 38, 974–980.

Sharp, I., Huang, F. and Patton, J. (2011). Visual error augmentation enhances learning in three dimensions. J NeuroEngineering Rehabil 8, 52.

Wang, Y. and Srinivasan, M. (2014). Stepping in the direction of the fall: the next foot placement can be predicted from current upper body state in steady-state walking. Biology Letters 10, 20140405.

Wu, A. R. and Kuo, A. D. (2016). Determinants of preferred ground clearance during swing phase of human walking. Journal of Experimental Biology jeb.137356.

Zelik, K. E., Huang, T.-W. P., Adamczyk, P. G. and Kuo, A. D. (2014). The role of series ankle elasticity in bipedal walking. Journal of Theoretical Biology 346, 75–85.

